# Randomness impacts specific priors building, visual exploration, and perception in object recognition

**DOI:** 10.1101/2023.09.26.559544

**Authors:** Cécile Gal, Ioana Țincaș, Vasile V. Moca, Andrei Ciuparu, Loredana E. Dan, Marie L. Smith, Teodora Gliga, Raul C. Mureșan

**Author notes:** Correspondence: Dr. Raul C. Mureşan, Transylvanian Institute of Neuroscience, Department of Experimental and Theoretical Neuroscience, Str. Ploiești 33, 400157 Cluj-Napoca, Romania. These authors contributed equally to the study (Cécile Gal, Ioana Țincaș).

## Abstract

Recognising objects is a vital skill on which humans heavily rely to respond quickly and adaptively to their environment. Yet, we lack understanding on the role visual information sampling plays in this process, and its relation to the individual’s priors. To bridge this gap, the eye-movements of 18 adult participants were recorded during a free-viewing object-recognition task using *Dots* stimuli (Moca et al., 2011). Participants viewed the stimuli in one of three orders: from most visible to least (*Descending*), least visible to most (*Ascending*) or in a randomised order (*Random*). This dictated the strength of their priors along the experiment. Adding to Moca et al.’s original finding that visibility order influenced participants’ recognition performance and visual exploration, we found that while orders allowing for stronger priors generally led participants to visually sample more informative locations, this was not the case of *Random* participants. Indeed, they appeared to behave naïvely, and their use of specific object-related priors seemed fully impaired, while they seemed to maintain the ability to use general task-related priors to guide their exploration. These findings have important implications for our understanding of perception, which appears markedly subjective even at the basic level of visual sampling and object perception.

## 2. Introduction

### 2.1. Background

Humans evolve in a highly complex visual world, in which visual scenes are composed of numerous objects, seen at different scales and angles, one often occluding another. Recognising objects under these conditions is a difficult task and yet every day we perform it seamlessly (Riesenhuber & Poggio, 2000). This process is extremely efficient and robust to challenging conditions, yet strikingly fast: human adults’ brain activity has been found to already present evidence of object recognition as fast as 150ms after stimulus presentation in electroencephalographic (EEG) recordings (Kirchner & Thorpe, 2006). This process is not static: it relies on active sampling (Findlay et al., 2003; Henderson, 2003). In fact, this seems to be a shared characteristic of all animals with a developed visual system, for whom visual behaviour is guided either through eye, head, or body movements (Land, 1999). These sequences of eye movements are not random: during the exploration of a scene, people preferably fixate informative locations, such as persons and objects (e.g. Yarbus, 1956). Humans’ visual behaviour is thus adapting not only to the varying physical properties of visual stimuli, like local contrast or spatial frequency (Ruddock et al., 1996; Zetzsche et al., 2000), but also to their individual circumstances, such as a goal sat by experimental instructions (Buswell, 1935; Yarbus, 1956; Castelhano et al., 2009) or relevant prior knowledge (Wu & Zhao, 2017).

Prior knowledge, or in short priors (Bernardo, 1979), is of particular interest because its influence on visual recognition suggests that people might not see the world in the same way given their past experience. In the context of visual search, priors have been proposed to be grouped into two main categories (Henderson, 2003): 1) *specific priors*, which relate to knowledge on the objects themselves and can be either short-term, acquired during the task, or long-term, acquired during the observer’s daily life; 2) *generic priors*, which relate to semantic and spatial knowledge providing general rules for how visual input is organised e.g., where to find a roof in a scene depicting a house. Such priors affect the recognition of objects by influencing the way observers sample visual information, but also their treatment of this information i.e., their perception. Priors can either help or hinder sampling and perception. For example, when a new category is learned, visually searching for an object that is part of this category among other objects is slower than for well-known categories, but it also results in less false alarms (Wu et al., 2013, 2016). Moreover, priors can also bias the perception of a stimulus: they can favour perceiving the same stimulus as previously seen in a hysteresis or attractive effect (Schwiedrzik et al., 2014; Snyder et al., 2015), or push the perception away from the previous stimulus in a repulsive or contrastive effect (Adams et al., 2013).

Thus, priors and eye movements central to the visual recognition process and can be studied in concert to characterise how one influences the other. Most of the work looking at priors and eye movements for object recognition has focused on visual search paradigms (see Henderson, 2003 for a review), which are particularly well-suited because they typically prompt extensive sequences of eye movements (Wolfe, 2020). Little is known about how priors influence eye movements in simpler tasks like single object recognition, because humans can habitually recognise single objects at a glance (Kirchner & Thorpe, 2006). Nonetheless, single objects have been shown to be explored extensively when they are difficult to perceive (Kietzmann & König, 2015; Moca et al., 2011), which offers new avenues for looking into the role of eye-movements, “*a window into the operation of the attentional system*” (Henderson, 2003, p. 498), for object recognition.

### 2.2. The Dots method

One method that can prompt intensified exploration of single objects through the alteration of visual stimuli is the *Dots methods* (Moca et al., 2011; Suzuki et al., 2018). In this type of paradigm, stimuli are composed of lattices of dots that are displaced away from their initial position towards contour-dense regions of an object, eventually revealing the contour of the object in regions where the dots come together. The deformation of the lattice is precisely calculated for each dot depending on the contour density of a source image combined with a global visibility variable, *g*. By varying the *g* value, one can vary the visibility of different stimuli in a highly controlled and comparable manner. Because the stimuli are all made of the same local elements, close comparisons are possible both between objects and between visibility levels. Dots stimuli are generated in a controlled manner, which bears the advantage of allowing to precisely quantify information content at any location in a stimulus. Two types of information can be extracted: 1) the *Local Dots Displacement (LDD)*, which represent the *physical information* present in the stimulus and varies with each level of visibility *g*; 2) the *Local Contour Density (LCD)*, which represents the *hidden information* from the source images and is independent of the *g*-level (see Moca et al., 2011 for more details). It is the latter that is used to generate the stimuli, but only the former is directly accessible to the participants. The two types of information are thus related but do not fully map onto each other. Reconstructing these values for locations fixated by participants allows to look into the way observers extract information through visual sampling. To make sense of Dots stimuli, participants have to rely on Gestalt principles, mainly proximity, grouping, and good continuation (Kubovy & Wagemans, 1995), and potentially to also rely on priors (Feldman, 1993, 1997, 2001). Following Henderson’s typology (2003), participants undergoing a Dots visual paradigm can be expected to use: 1) *specific* priors related to the particular *objects* of the stimulus-set, which they can both build as they recognise the objects and withdraw from their daily-life experience; and 2) *generic* priors that provide *contextual* rules about how the stimuli are presented within the task e.g., their order and composition, acquired throughout the experiment.

The Dots method was introduced by Moca et al. (2011), who presented participants with 50 objects of 7 visibility levels *g*, in seven blocks of incremental *g* (0.00-0.30). This was a way to manipulate participants’ access to information and thereby their ability to build priors right from the start of the experiment. Participants in an *Ascending* group experienced a block-by-block increase of the visibility level *g*, while participants in a *Descending* group saw blocks of decreasing visibility. Moca et al. showed that prior access to information influenced participants’ perception, i.e., their ability to accurately recognise the objects, but also their visual exploration of the stimuli, i.e., their ability to sample informative points. The *Descending* group who could build the strongest priors from the start both recognised stimuli more accurately and sampled more informative locations, especially at lower visibility levels, compared the *Ascending* participants.

### 2.3. Current study

The current study extends the work of Moca et al. (2011), whose data was revisited and reanalysed, alongside a further group of *Random* participants. This additional group of participants viewed the same stimuli as the former *Ascending* and *Descending* participants, with each of the 50 objects similarly presented once per block, but this time *g*-levels were randomised across the seven blocks. This *Random* group of participants was thereby given an intermediate access to information compared to the *Ascending* and *Descending* participants and was expected to build priors, explore, and perform at an intermediate level between the two former groups. Additionally, here we introduce a simple model (see **Figure 1**), to provide more insight into the recognition process. The model sets out to account for changes in performance at three main levels of visibility (left to right: low, medium, and high *g*) in terms of 1) the *available information* (top row); 2-3) the *task-general* (generic) and *object-specific* (specific) *priors* (respectively, second and third row); and 4) the *performance* (bottom row), for each of the three groups. An additional, purely theoretical naïve group who is not building any priors but experiencing varying levels of information availability throughout the *g*-levels, was also added to the model as a reference point for purely stimulus (*g*)-related, prior-unrelated changes in performance.

**Figure 1.**
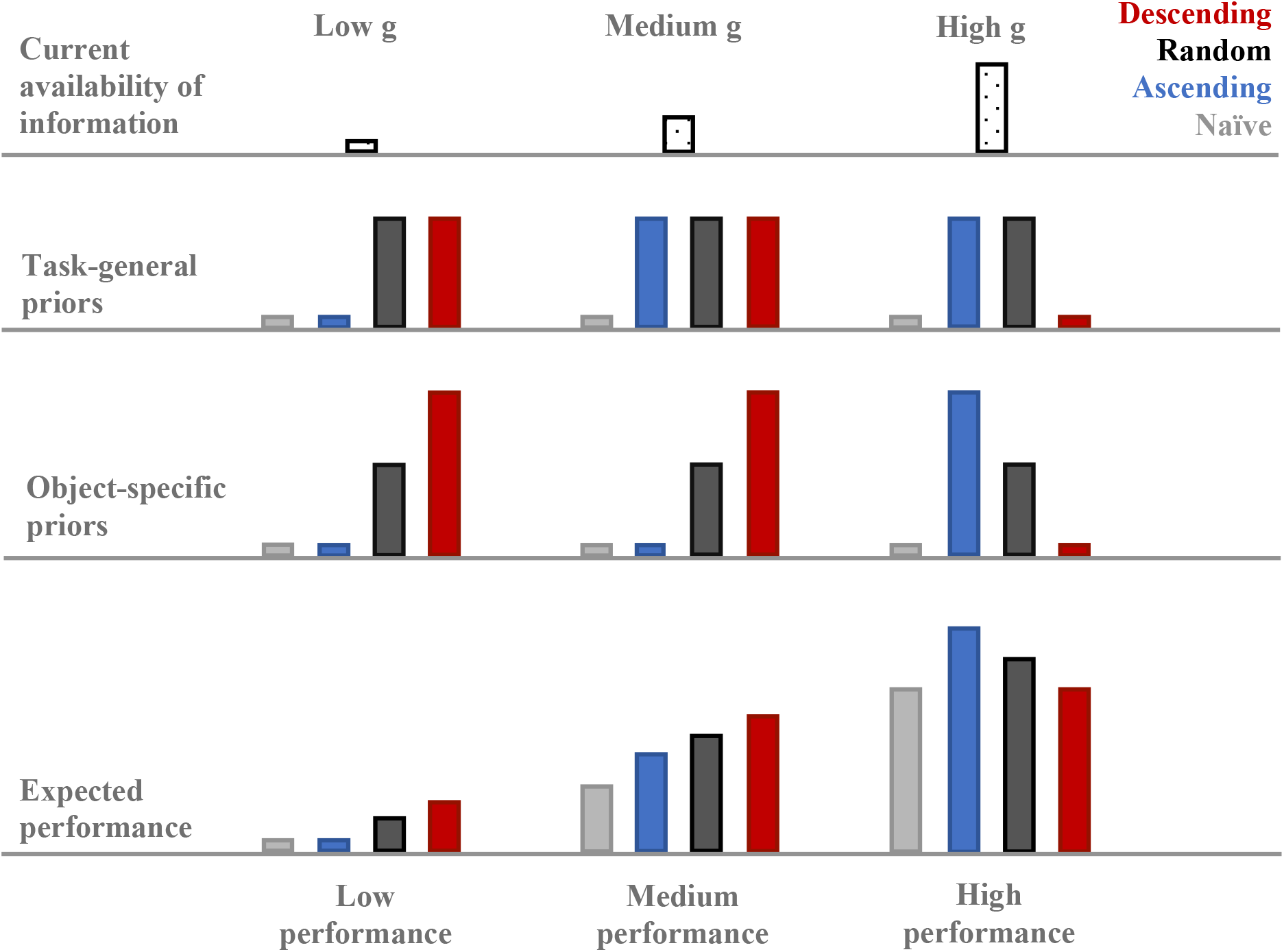
Qualitative model of participants’ access to information (top row), task-general (middle-top) and object-specific (middle-low) priors, as well as expected performance (bottom) along 3 generic *g*-levels: low (left column), medium (middle column), and high (right column). *Descending* participants are depicted in red, *Ascending* ones in blue, and *Random* ones in black. Grey bars correspond to a hypothetical group of naïve participants building no priors. Higher bars signify higher values.

We predict, as seen for the two original groups in Moca et al. (2011), that all three groups will perform better with increasing *g*, and that each group’s performance relative to the other groups will depend on the strength of their priors at each *g*. We expect the three groups to build task-general priors equally well depending on their mere number of trials, as reflected by symmetrical changes along *g*-levels for the *Ascending* and *Descending* groups, and constant, intermediate general priors at all *g*-levels for the *Random* group (second row). By contrast, the object-specific priors are thought to be built according to present and past information availability: *Descending* participants build strong priors quickly, while *Ascending* ones only build limited priors from medium *g* onward, and *Random* participants’ priors should be moderately strong all throughout (third row). Thus, we predict that *Descending* participants will perform best at low and medium *g*, while *Ascending* participants will dominate at high *g*, and the *Random* group will remain intermediate all throughout. Conversely, we expect these groups to match the theoretical naïve group when they start the experiment: at high *g* for the Descending group and low *g* for the Ascending group. We expect all these effects to be reflected in participants’ performance (bottom row), both in terms of recognition accuracy and visual exploration informativeness (LCD and LDD).

## 3. Results and comparison to model

### 3.1. Recognition accuracy

Participants’ recognition accuracy (answered “Seen” and correctly recognised; **Figure 2**) was analysed for the three groups of *Ascending*, *Descending,* and *Random* participants and for the seven *g*-levels in a 3*7 repeated measures analysis of variance (RM-ANOVA) using a Huynh-Feldt correction for non-sphericity. To note, unequal variances were also found for most *g*-levels, suggesting that the homogeneity of the participants’ responses varied as a function of *g*. This was expected, as the low *g*-levels corresponded to no or very little visibility, causing a floor effect, while the highest *g*-levels created a ceiling effect, and the levels in between were expected to be particularly affected by the groups’ difference in priors’ strength. No correction could be applied to correct for this violation as well as the violation of sphericity, however since the groups had the same sample size, it was considered appropriate to still conduct ANOVAs (Weerahandi, 1995). Indeed, the violation of the equal variance assumption yields an increased likelihood of type II error or false negatives compared to equal variances, while the type I errors or false positives remained comparable, resulting in more stringent tests such that positive results remained highly reliable.

**Figure 2.**
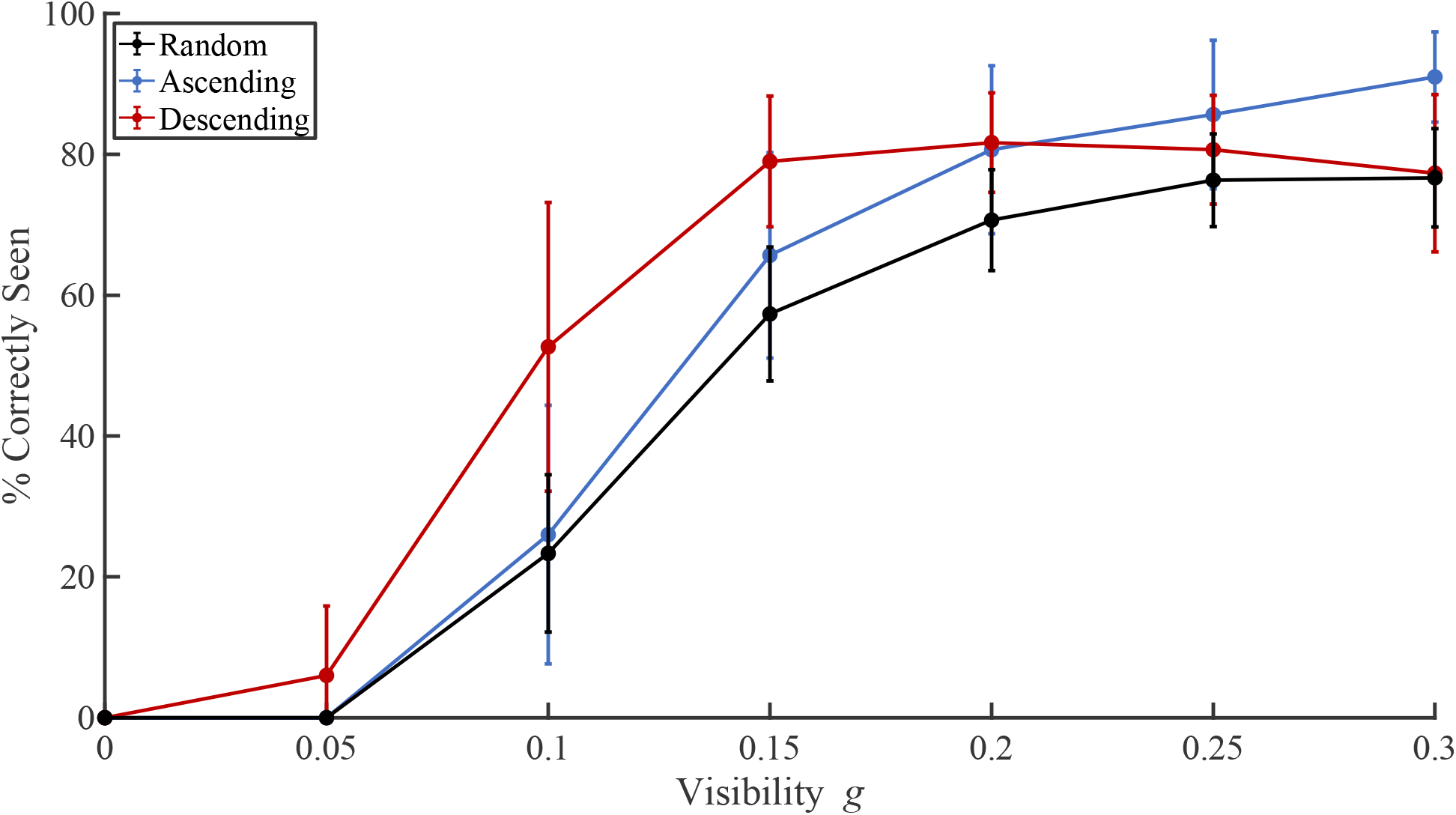
Recognition accuracy as a function of *g* and for each group. Error bars represent the standard deviation.

There was a strong effect of *g* on recognition accuracy (F(2.759,41.384) = 389.526, p < 0.001, η^2^ = 0.887): all participants appeared to recognise the objects increasingly well for larger visibility levels. There was no evidence that the different groups recognised objects overall better or worse than the others, as reflected by no significant *group* effect (F(2,15) = 2.617, p = 0.106, η^2^ = 0.014). However, participants’ response accuracy changed with the visibility *g* in a way that depended on their group, as reflected by a significant *g*group* effect (F(5.518, 41.384) = 5.230, p < 0.001, η^2^ = 0.024). Indeed, as seen in Moca et al. (2011) and in accordance with the model (**Figure 1**), the *Ascending* participants appeared to dominate at high *g*, while the *Descending* group was more accurate at middle and low *g*, which in both cases corresponded to the levels when they had the strongest priors. On the other hand, the *Random* group did not appear to follow the model’s predictions: their recognition accuracy remained comparable to the group with weak priors and lower than the group with strong priors all along, never reaching the intermediate level predicted (contrasts at medium *g*: t(1,21.01) = 1.398 and p = 0.177 for *Random* vs. *Ascending*, t(1,21.012) = 4.127 and p < 0.001 for *Random* vs. *Descending*; at high *g*: t(1,26.81) = 2.216 and p = 0.035 for *Random* vs. *Ascending*, t(1,26.81) = 0.468 and p = 0.644 for *Random* vs. *Descending*). This suggests a poorer building of priors than expected for this group, who appeared to be behaving like the model’s theoretical naïve group.

#### 3.1.1. Recognition accuracy as a function of blocks

To test this idea, post-hoc analyses were conducted on the *Random* group’s response accuracy by *block*, rather than by *g*, which allowed us to look for a performance improvement over time (**Figure 3**). We projected no effect this time, since it would suggest that these participants were building and using priors over time as they accumulated information, as opposed to behaving naïvely as we just proposed. This analysis was not performed on the other groups as their block order was the same as (or the reverse of) the *g* order, which did not allow to decouple visibility and learning effects. The data was analysed in a Bayesian RM-ANOVA such that evidence for the null hypothesis could be examined. We found decisive evidence (BF > 100) for the alternative hypothesis supporting an effect of block: BF_10_ = 256.522. However, this effect seemed to be driven by the change between block 1 and 2, as shown by a dramatic drop to an anecdotal value (1/3 < BF < 1) in favour of the null hypothesis when not including block 1 in the analysis: BF_10_ = 0.995. This implies that the *Random* participants did improve their performance and built priors, but only during the first block and not along the whole experiment as was initially expected. This was taken as reflecting the accumulation of task-general priors only. Indeed, in only a few trials participants could quickly acquire good knowledge on the general properties of the stimuli with which they were presented, especially as they first performed 7 practice trials (see Methods section), which made successive trials uninformative regarding the building of this type of general priors. By contrast, accumulating object-specific priors demanded several blocks since many of the stimuli that they saw in the first block were sub-recognition-threshold and did not enable them to build priors yet.

**Figure 3.**
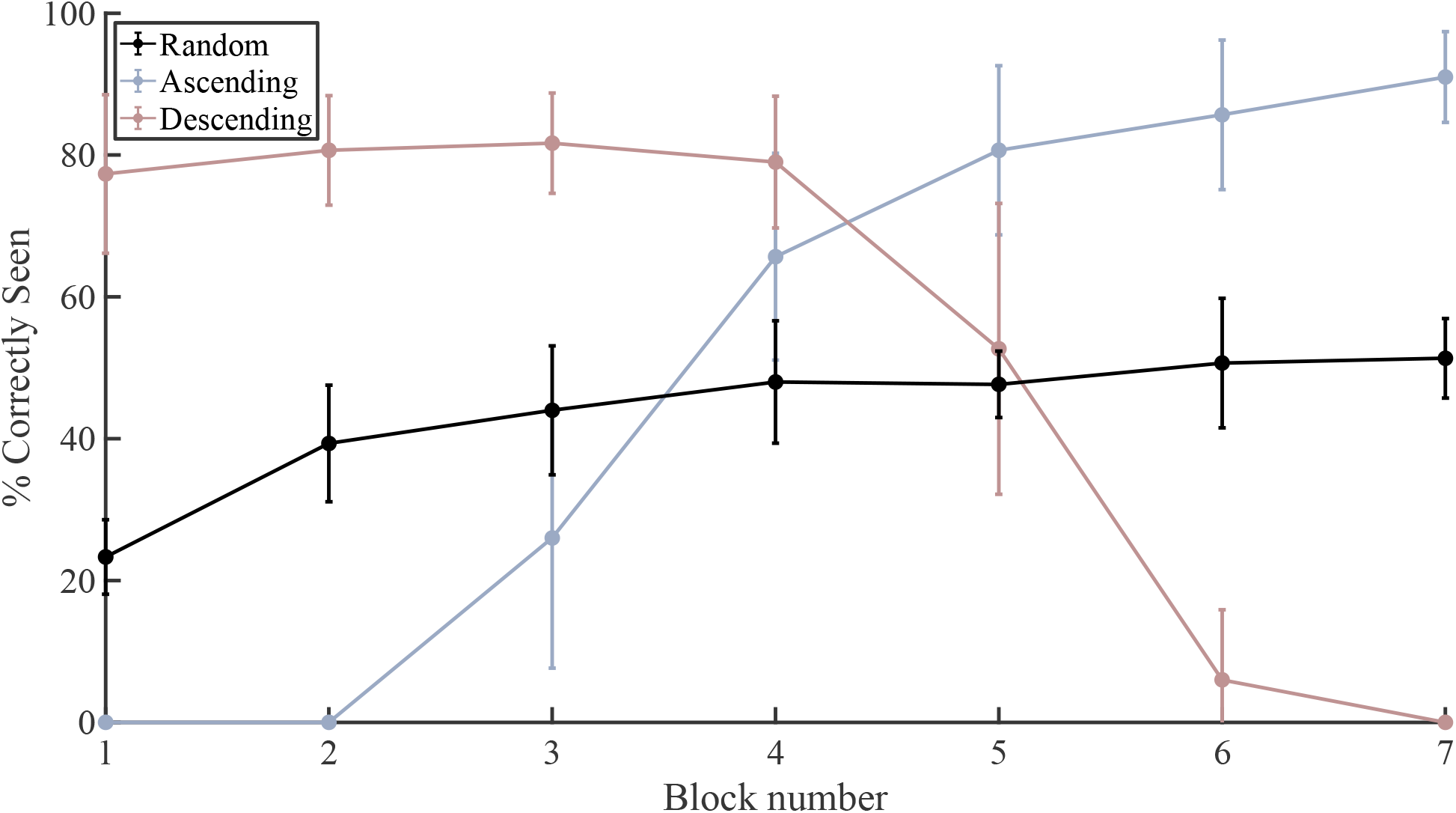
Recognition accuracy as a function of *block* for the *Random* group (black). The values for the *Ascending* and *Descending* group, while not analysed, are still depicted for reference in shaded blue and red, respectively. Error bars represent the standard deviation.

Hence, the *Random* group’s performance was re-computed not taking block 1 into account (**Figure 4**) to test whether decoupling the *Random* participants’ learning effect at the start of the experiment from their performance over different *g*-levels at later stages in the task enhanced their performance relative to the other groups. We found that their performance curve shifted up towards a more accurate recognition, but the change was minimal, and the group remained at a comparable or lower level than the weak-prior groups, never reaching the expected intermediate level (contrasts at medium *g*: t(1,46.511) = 0.635 and p = 0.517 for *Random* vs. *Ascending*, t(1,46,511) = 2,741 and p = 0.009 for *Random* vs. *Descending*; at high *g*: t(1,46.511) = 2.114 and p = 0.040 for *Random* vs. *Ascending*, t(1,46.511) = −0.026 and p = 0.979 for *Random* vs. *Descending*).

**Figure 4.**
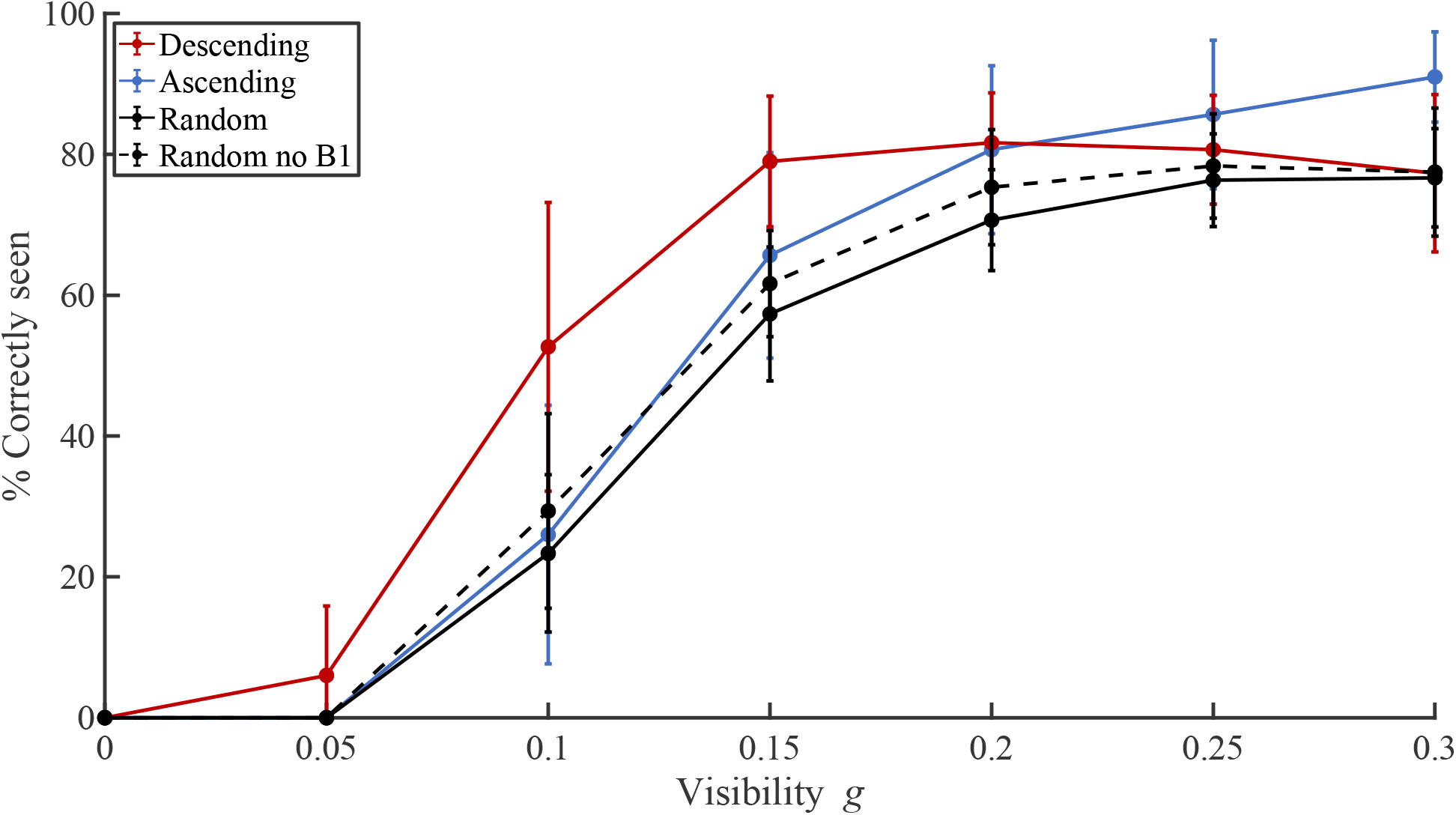
Recognition accuracy as a function of *g* and for each group, including the *Random* group without block 1 (black dashed lines). Error bars represent the standard deviation.

### 3.2. Visual behaviour

In a second set of analyses, we investigated whether the effects of visibility (*g*-level) and priors (group: *Ascending*, *Descending* or *Random*) on recognition performance could be linked to differences in participants’ visual information sampling. To this end, participants’ eye-movements were recorded using eye-tracking, and the information content at the locations fixated by the participants was analysed. Using the *Dots method* (Moca et al., 2011), we extracted measures of *available* information, or Local Dots Displacement (LDD), and *hidden* information, or Local Contour Density (LCD) at each point fixated. For each type of information, two measures were extracted on a trial-by-trial basis: the average LDD or LCD value across all fixations made within a trial, and the total information sampled within a trial (the sum across the trial’s fixations). The former measure is considered to capture participants’ strategy with regards to sampling informative locations, while the latter reflects the amount of information that the participants needed in order to reach a decision about the identity of the object in a given trial. Similarly to the first analysis, this data was explored as a function of participant group (three levels) and *g* (seven levels) in a 3*7 RM-ANOVA with a Huynh-Feldt correction for sphericity violation while unequal variances remained uncorrected as justified before (Weerahandi, 1995).

#### 3.2.1. Local Dots Displacement (LDD)

In terms of *available* information (LDD, **Figure 5**), we found a strong, positive effect of *g* both on the average and the total LDD accessed by participants (respectively: F(2.753,41.297) = 1088.508, p < 0.001, η^2^ = 0.981; and F(2.159,32.379) = 37.768, p < 0.001, η^2^ = 0.449). For the average LDD, this was not accompanied by any effect of group (F(2,15) = 0.834, p =0.454, η^2^ = 0.0004) or g*group (F(5.506,41.297) = 0.866, p = 0.520, η^2^ = 0.002): this showed a relatively unbiased, prior-independent sampling of the stimulus space by participants in terms of *available* (physical) information. In terms of total LDD however, we found a significant interaction effect of g*group (F(4.317,32.379) = 5.494), p < 0.001, η^2^ = 0.131) but no group effect (F(2,15) = 0.251, p = 0.781, η^2^ = 0.008). This suggests that while all participants similarly sampled more information at each fixation on average when information availability *g* increased, the amount of information that they needed to reach their decision given the *g*-level depended on their group. Here the model’s predictions are particularly helpful to explain the seemingly complex patterns that Moca et al. (2011) had documented. Indeed, the *Ascending* group needed the least amount of information compared to the other groups at high *g* when the model predicts that their priors were the strongest (contrast with both other groups at *g* = 0.30: t(1,42.495) = 7.644, p < 0.001), while the *Descending* group needed the highest amount of information at high *g* when they were naïve (contrast with both other groups at *g* = 0.30: t(1,42.495) = 0.799, p = 0.429) and the lowest at middle *g* when they had the strongest priors (contrast with both other groups at *g* = 0.15: t(1,42.495) = 4.352, p < 0.001). The *Random* group appeared intermediate at both medium and high *g*, which was compatible with our model, but needed the least amount of information at low *g*, which we predicted would rather be the case for the *Descending* group (contrast with both other groups at *g* = 0.05: t(1,4) = 2.341, p = 0.024). Interestingly, the *Random* group’s need for *physical* information before reaching a decision scaled linearly and did not match their relative recognition performance (which was shown earlier to be lower or similar than both groups all along and never intermediate or higher). This possibly reflects an overall lower motivation.

**Figure 5.**
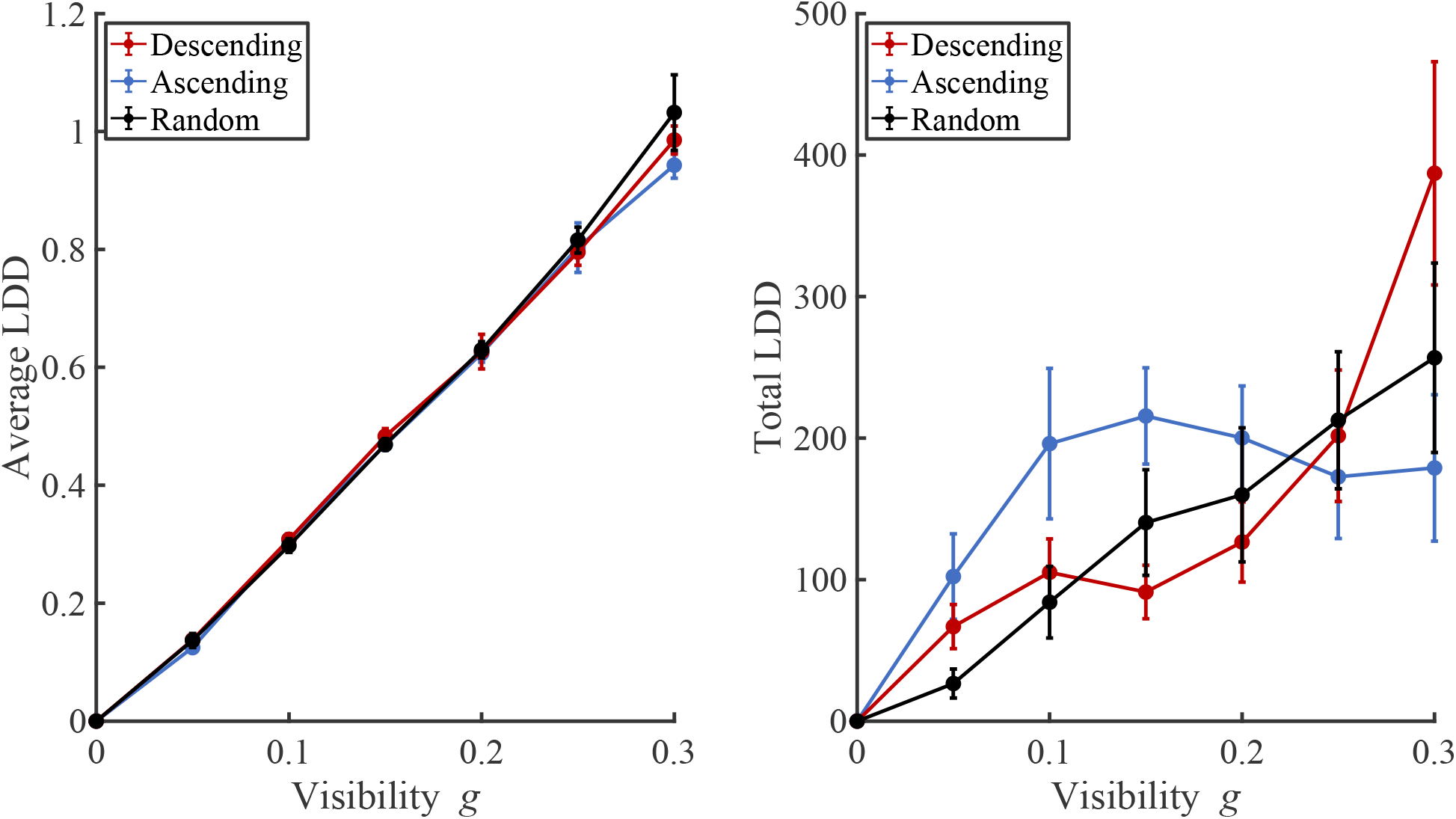
Average of the trials average (left) and total (right) Local Dots Displacement (LDD) as a function of *g* and for each group. Error bars represent the standard deviation.

As for the former analyses, we looked at potential learning effects along blocks in the *Random* group (**Figure 6**). We found no effect of block for the average LDD (F(6, 96) = 1.623, p = 0.149, η^2^ = 0.135), which was compatible with the former result that the average *physical* LDD sampled by participants appeared to be more a function of the visibility *g* than of their priors’ strength. On the other hand, we found a significant block effect for the total LDD (F(6,96) = 2.488, p = 0.028, η^2^ = 0.135), which appeared to decrease with blocks. This suggests that as they progressed through trials, *Random* participants either needed less information to reach a decision or became less motivated to explore. To note, this general decrease appeared marked by an increase after they started and as they finished the experiment, which was particularly compatible with a motivational perspective as participants were notified of how many blocks were left at each inter-block interval, while their performance did not seem to follow the same pattern.

**Figure 6.**
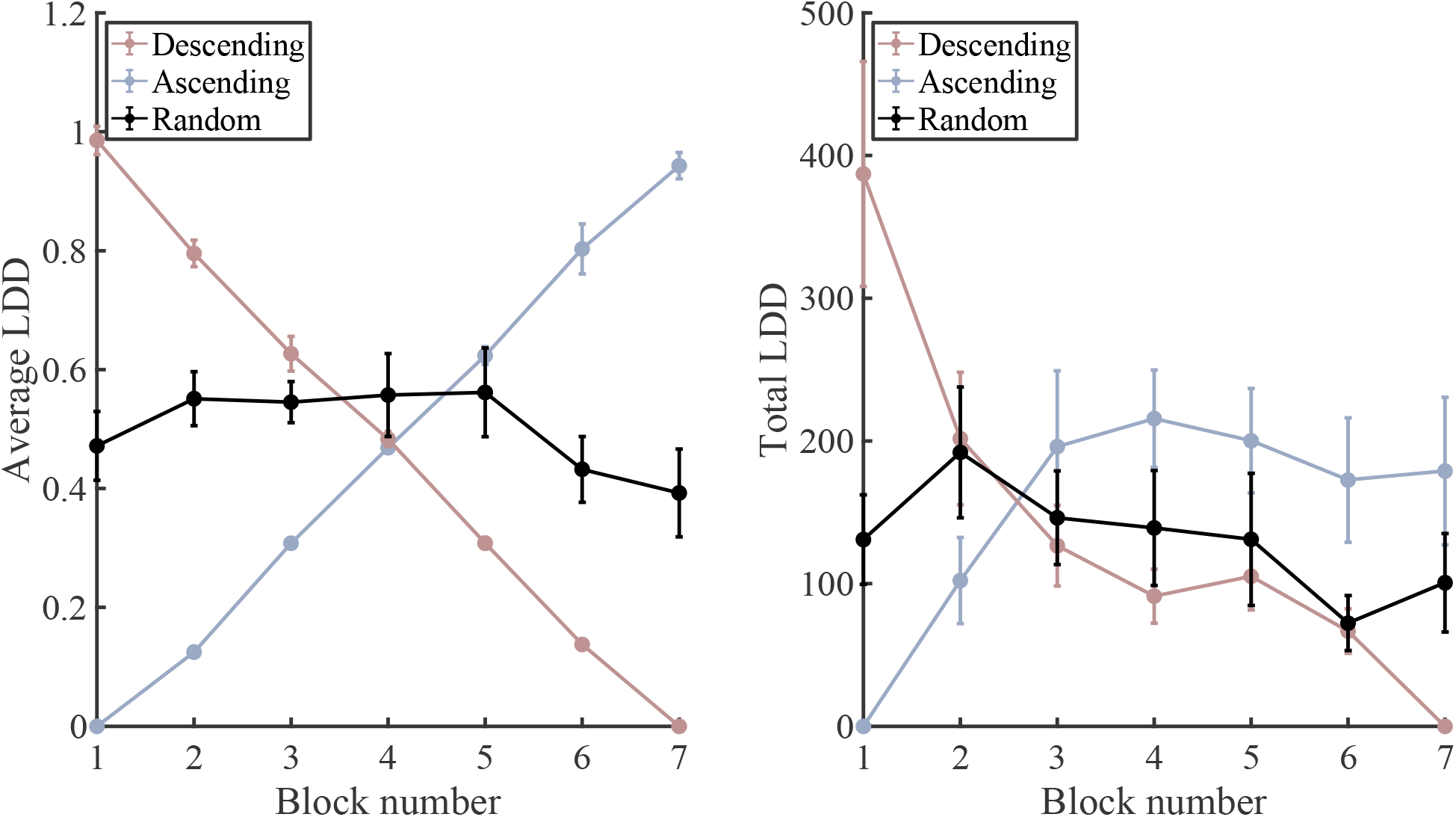
Average (left) and total (right) Local Dots Displacement (LDD) as a function of *block* for the *Random* group (in black). The values for the *Ascending* and *Descending* group, while not analysed, are still depicted for reference in shaded blue and red, respectively. Error bars represent the standard deviation.

#### 3.2.2. Local Contour Density (LCD)

In terms of *hidden* information, or LCD (**Figure 7**), we found a significant effect of *g* on the average LCD (F(4.747, 71.204) = 51.184, p < 0.001, η^2^ = 0.585), coupled with a significant interaction effect of g*group (F(9.494, 71.204) = 2.236, p = 0.027, η^2^ = 0.051) and no significant effect of group (F(2,15) = 0.789, p = 0.472, η^2^ = 0.018). This suggests that participants were becoming more efficient at sampling *hidden* information (local contour from the original image) as visibility increased, in a way that depended on their priors. Indeed, as predicted by the model and similarly to the total LDD described earlier, the *Ascending* group accessed locations with the highest average LCD at high *g* (contrast with both other groups at *g* = 0.30: t(1,50.940) = 10.815, p < 0.001), while it was the *Descending* group who sampled locations with the highest LCD at middle *g* (contrast with both other groups at *g* = 0.15: t(1,50.940) = 10.442, p < 0.001). The *Random* group remained generally below both the other groups all throughout (contrast with both other groups pooling all *g*-levels: t(1,15.000) = 15.510, p < 0.001), similarly to their performance in terms of recognition accuracy and resembling the behaviour of the naïve group. There was an exception for the first *g*-level, when there was a surge in the *Random* group’s average LCD sampled and the group actually reached a level comparable to the *Descending* group (contrast at *g* = 0.00: t(1, 50.940) = −0.799, p = 0.428) and higher than the *Ascending* group (contrast at *g* = 0.00: t(1, 50.940) = - 2,626, p = 0.011), at odds with the idea that this group did not build priors. Interestingly, this *g*-level actually did not contain any *physical* LDD information, so any of the *hidden* LCD information sampled at this point was guided not by the physically available information or by object-specific priors, but by a combination of chance and task-general priors (knowledge about the location and extent of objects in the stimulus images). Thus, this result at the lowest *g*-level again implies that the *Random* group’s impaired ability to guide their behaviour with priors might have been limited to object-specific priors only, while their ability to build and use task-general priors was maintained.

**Figure 7.**
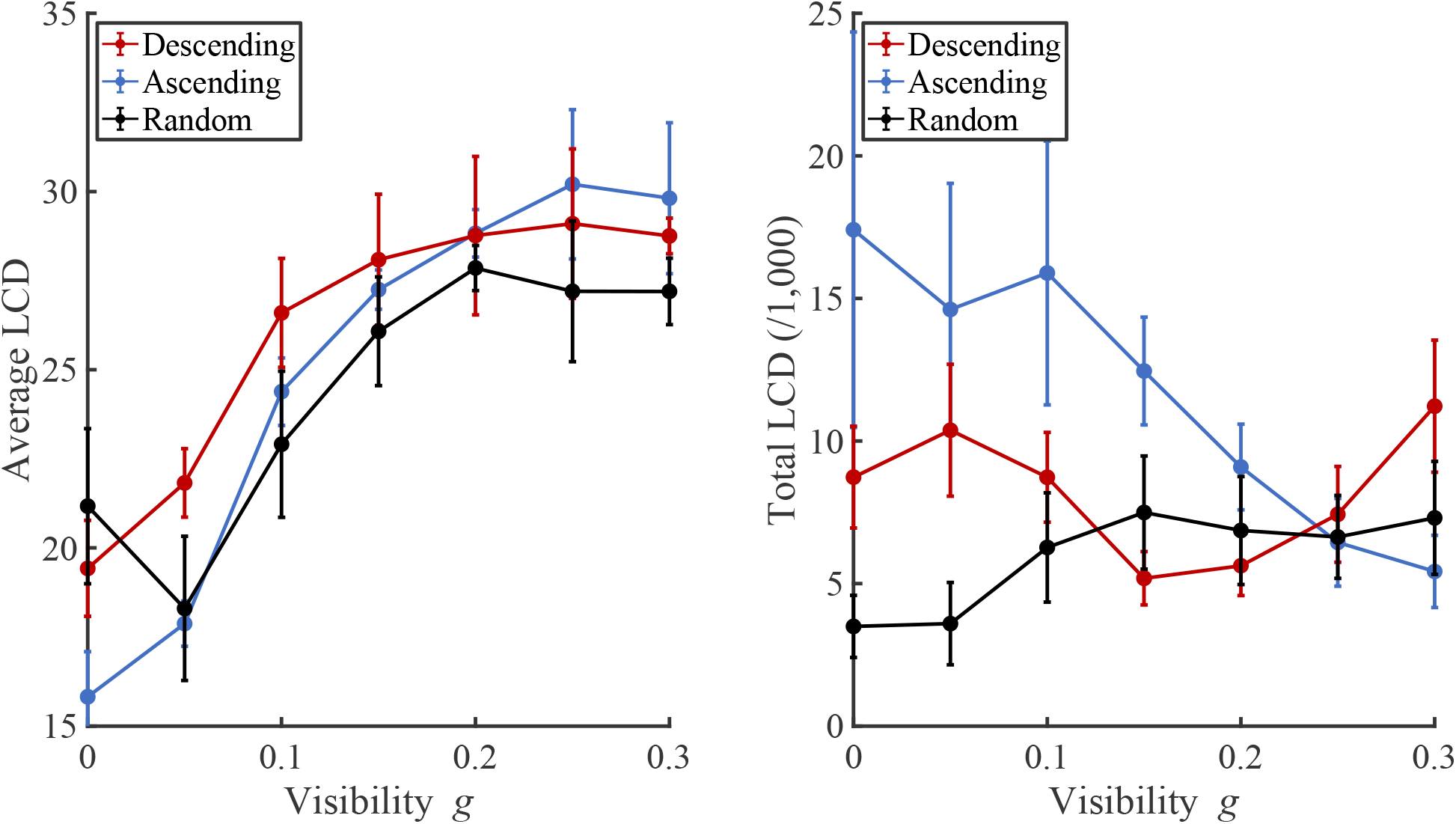
Average (left) and total (right) Local Contour Density (LCD) as a function of *g* and for each group. Error bars represent the standard deviation.

The effect of *g* on the total LCD was not significant, although close to significance threshold (F(2.001, 30.009) = 2.174, p = 0.053, η^2^ = 0.038) and associated with a significant g*group effect (F(4.001,30.009) = 4.922, p = 0.004, η^2^ = 0.173) and no significant group effect (F2,15) = 2.542, p = 0.112, η^2^ = 0.133). This suggests that the amount of *hidden* LCD information that participants needed to integrate in order to reach a decision changed as a function of *g* in a group-dependent manner. Qualitatively, the changes in total LCD with *g* appeared to follow non-linear and group-specific shapes. The *Ascending* group sampled decreasing amounts of total LCD with *g* (simple main effect of *g*: F(1,6) = 3.814, p = 0.006), as both their priors and access to information increased block by block, and resulting in them sampling the least total LCD at the highest *g*-level (contrast with both other groups at *g* = 0.30: t(1,38.945) = 3.268, p = 0.002).

The *Descending* group followed a more complex pattern: their total LCD sampled also decreased with blocks at the start of the experiment, reflected by a decrease from high to middle *g* as the group built priors, resulting in this group accessing the smallest amount of total LCD at middle *g* (contrast with both other groups at *g* = 0.15: t(1,38.945) = 3.682, p < 0.001). As the information became scarcer in the next blocks, *the Descending* group sampled more total LCD as *g* continued to decrease, possibly to compensate for the gradual vanishing of information from the stimuli. The *Random* group’s total LCD was stable at middle and high *g* but was lower for low *g*-levels, when they explored the least of all groups (contrast with both other groups at *g* = 0.00: t(1,38,945) = 5,645, p < 0.001). This indicates a premature cessation of exploration in the *Random* group, compatible with the idea of a curb in motivation, especially when compared to the *Descending* participants increased effort with decreasing *g*.

Once more, we looked at these measures as a function of block for the *Random* group (**Figure 8**). We found no effect of block on average LCD (F(6,96) = 1.741, p = 0.120, η^2^ = 0.098) nor total LCD (F(6,96) = 1.766, p = 0.114, η^2^ = 0.099), suggesting that *Random* participants did not learn to sample more *hidden* information over the experiment. Once more, the fact that this group did access less total *physical* but not *hidden* information suggests that their building of task-general priors remained functional while their object-specific prior building was impaired.

**Figure 8.**
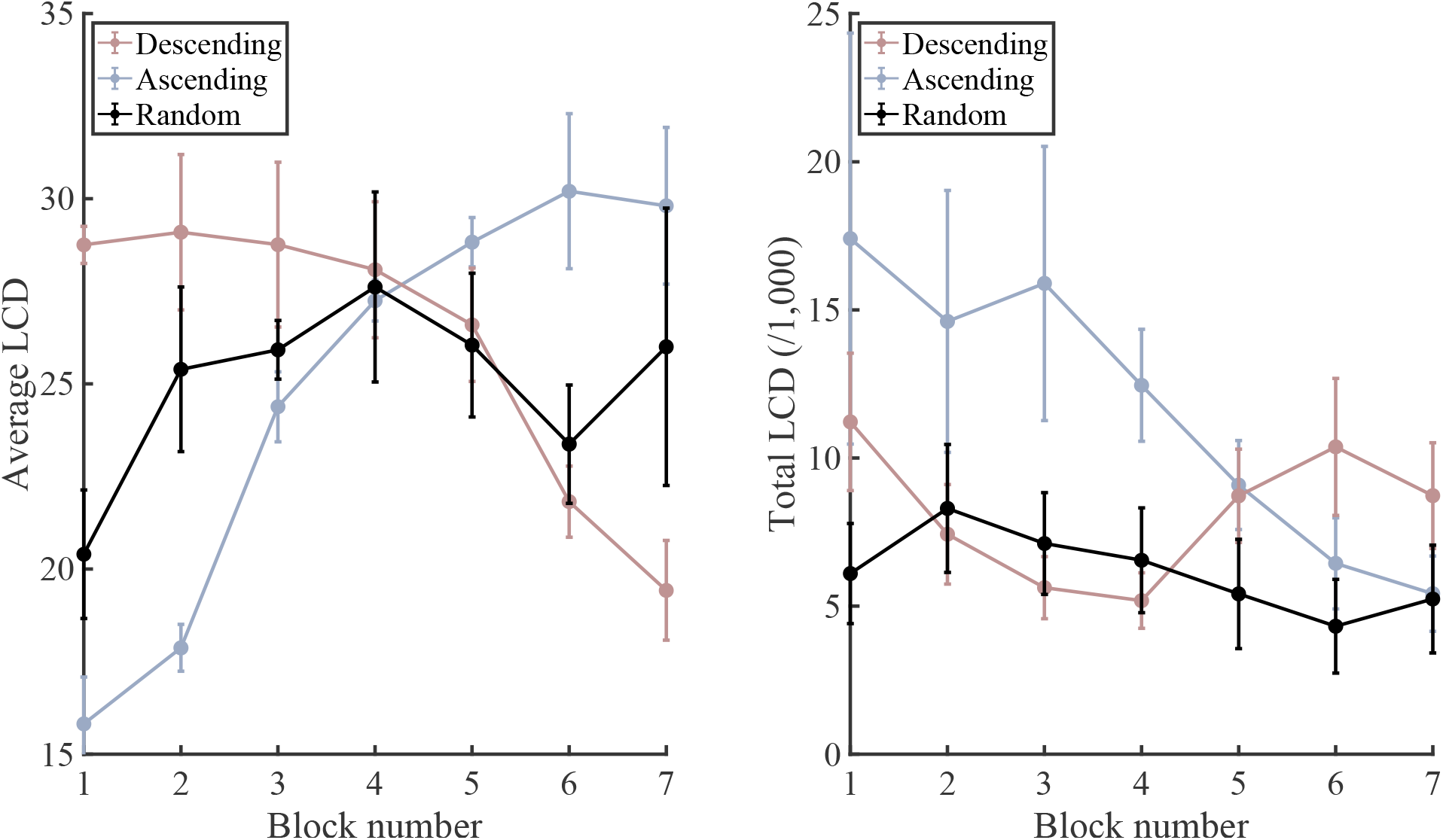
Average (left) and total (right) Local Contour Density (LCD) as a function of *block* for the *Random* group (in black). The values for the *Ascending* and *Descending* group, while not analysed, are still depicted for reference in shaded blue and red, respectively. Error bars represent the standard deviation.

### 3.3. Lateralisation of the fixations

Because the *Dots* stimuli require participants to rely on Gestalt principles of proximity and good continuity, focusing exactly on the informative points is not necessarily the most efficient strategy for exploring these stimuli. Indeed, grouping dots together into a meaningful contour can be easier when the dots at hand are besides the gaze’s focus point, where the peripheral vision blurs the points together which can then appear linked in a contour. Lateralising the fixations to the right of the location of interest has been shown to be helpful in stimuli made of dots, because they reached the right cerebral hemisphere that is more efficient in identifying meaningful patterns (Brugger & Regard, 1995; Dan et al., 2022). Thus, based on the evidence that the Random group, contrary to our model’s predictions, did not sample as much information as they could afford to given their prior access to information, we performed a last post-hoc analysis looking into the lateralization of the participants’ fixations. Indeed, participants might have been relying more on lateralisation when either their priors or their access to information was limited: lateralising could have been a compensation strategy used by participants, resulting in them apparently sampling less information, while actually being able to find the information and precisely gaze next to it.

Lateralisation (**Figure 9**) appeared to significantly shift from right to left both along *g* levels (F(2.699,40.484) = 2.954, p = 0.049, η^2^ = 0.055, Greenhouse-Geisser corrected) and *blocks* (F(2.740,38.356) = 4.118, p = 0.0115, η^2^ = 0.093, Greenhouse-Geisser corrected). This was not associated with a *g*\*group (F(5.398, 40.484) = 0.232), p = 0.400, η^2^ = 0.040, Greenhouse-Geisser corrected) or a *block*group* (F(5.479, 38.356) = 0.678, p = 0.656, η^2^ = 0.031, Greenhouse-Geisser corrected) effect, suggesting that participants’ change in lateralisation was affected in a similar way across groups both by the visibility of the stimuli (*g*) and by their experience with the stimulus set (*block*). However, there was also a significant effect of *group* (F(2,15) = 5.143, p = 0.020, η^2^ = 0.254), suggesting that although all participants were similarly affected by *g* and *block*, their overall degree of lateralisation was not the same depending on the order in which they saw the stimuli (*group*). Indeed, *Random* participants appeared to lateralise generally more than the other groups, although this effect only came close to significance (contrast between the *Random* group and both other groups: t(1,15.000) = - 2.107, p = 0.052).

**Figure 9.**
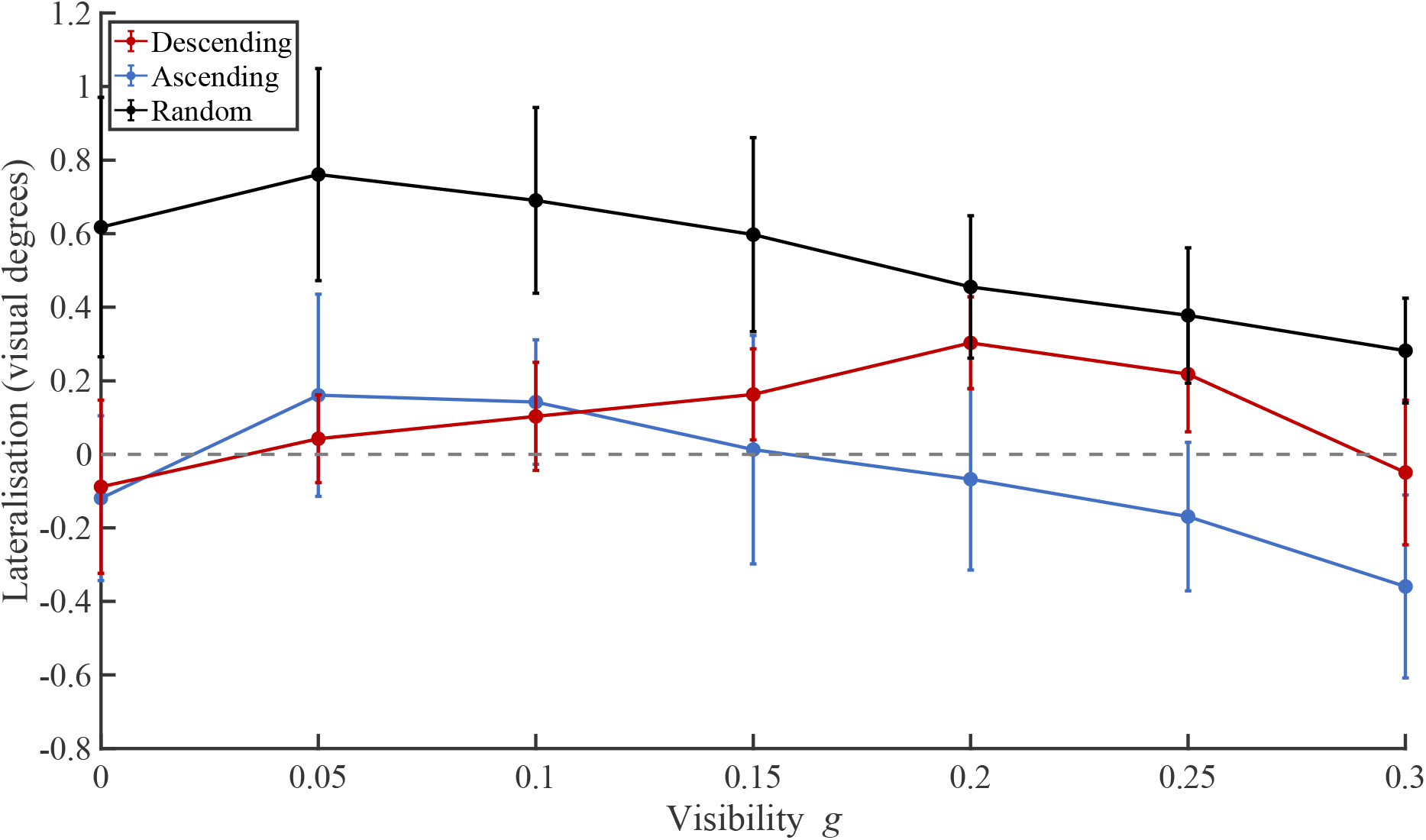
Fixation lateralisation as a function of *g* and for each group. Error bars represent the standard deviation. Values above zero represent a right lateralisation and values below zero, a left lateralisation.

**Figure 10.**
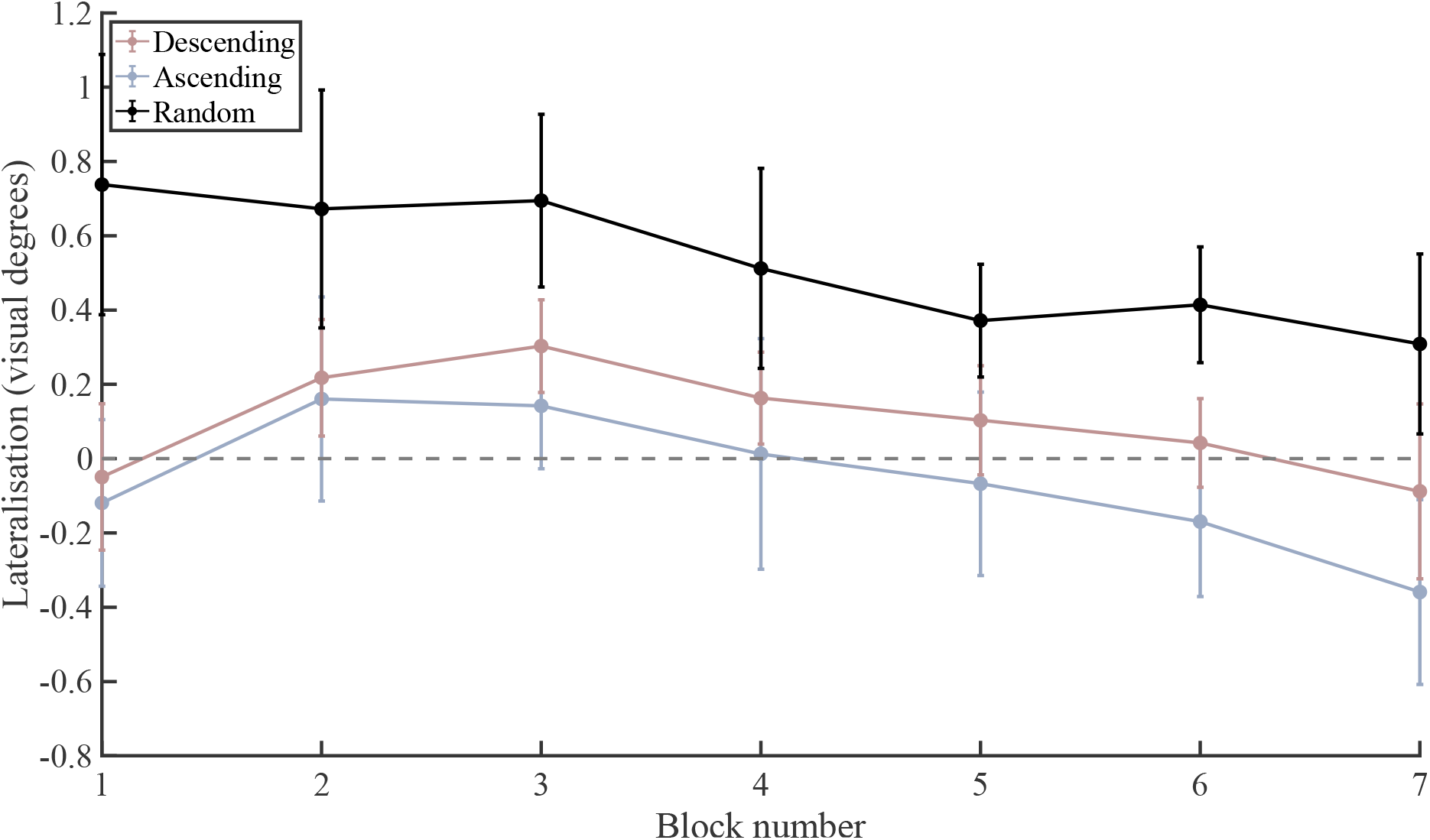
Fixation lateralisation as a function as a function of *block* for (*Random* group in black). The values for the *Ascending* and *Descending* group, while not analysed, are still depicted for reference in shaded blue and red, respectively. Error bars represent the standard deviation. Values above zero represent a right lateralisation and values below zero, a left lateralisation.

## 4. General discussion

We showed that participants’ access to information, both in-the-moment (visibility *g*) and preceding (priors) the stimulus at hand, influenced their prior building, visual exploration, and perception. Indeed, by presenting stimuli of objects in blocks of either *Ascending*, *Descending*, or *Random* visibility order, we controlled participants’ ability to build priors, which allowed us to compare recognition performance and exploration between different information-access scenarios. In line with Moca et al. (2011), all participants were better at both recognising and exploring the informative locations of stimuli as visibility (*g*) increased. However, each group’s performance relative to the others changed along *g*, which we showed was well explained by a simple model (**Figure 1**) combining the in-the-moment availability of information (*g*) with the previous access to information (priors) for the *Ascending* and *Descending* groups. Indeed, the *Ascending* participants performed the best at the highest *g*-levels, when they had already gone through all the lower *g*-levels and held the strongest priors of all groups. They both recognised objects more accurately and explored more informative locations, while needing the smallest amount of information to reach a decision about objects identity. At medium *g*-levels, this was true for the *Descending* group, who had just gone through blocks of the most visible levels and held the strongest priors of all groups at this point.

The *Random* group, which was not analysed in the original Moca et al. (2011) study, was found not to perform according to our model. *Random* participants appeared to both explore and recognise objects the least well of all tree groups at all *g*-levels, despite their access to information being intermediate between the two former groups, which according to the model should have led to an intermediate performance. This result suggests that randomness impairs *Random* participants’ ability to build and use priors to guide their recognition and exploration of the stimuli. Interestingly, while it appears that *Random* participants’ *object-specific* priors were hindered and could not guide their exploration and recognition of objects, several pieces of evidence suggest that this was not the case of their *task-general* priors. Indeed, *Random* participants’ performance improved over the first block only, which is not compatible with the accumulation of object-specific information over several blocks, as more and more visible stimuli keep being presented over time, but is compatible with the evaluation of task-general statistics which do not change from a block or a stimulus to another and can be learned quickly from the start. Moreover, *Random* participants appeared to perform as well as the high-prior *Descending* group and better than the naïve *Ascending* group at the lowest level *g* = 0.00. At this level, there was no physically *available* information, such that participants could only be guided by their task-general priors to access areas that were statistically informative. Accordingly, we propose an updated version of our model accounting for this limitation in performance and lack of specific prior building in the *Random* group (**Figure 11**).

**Figure 11.**
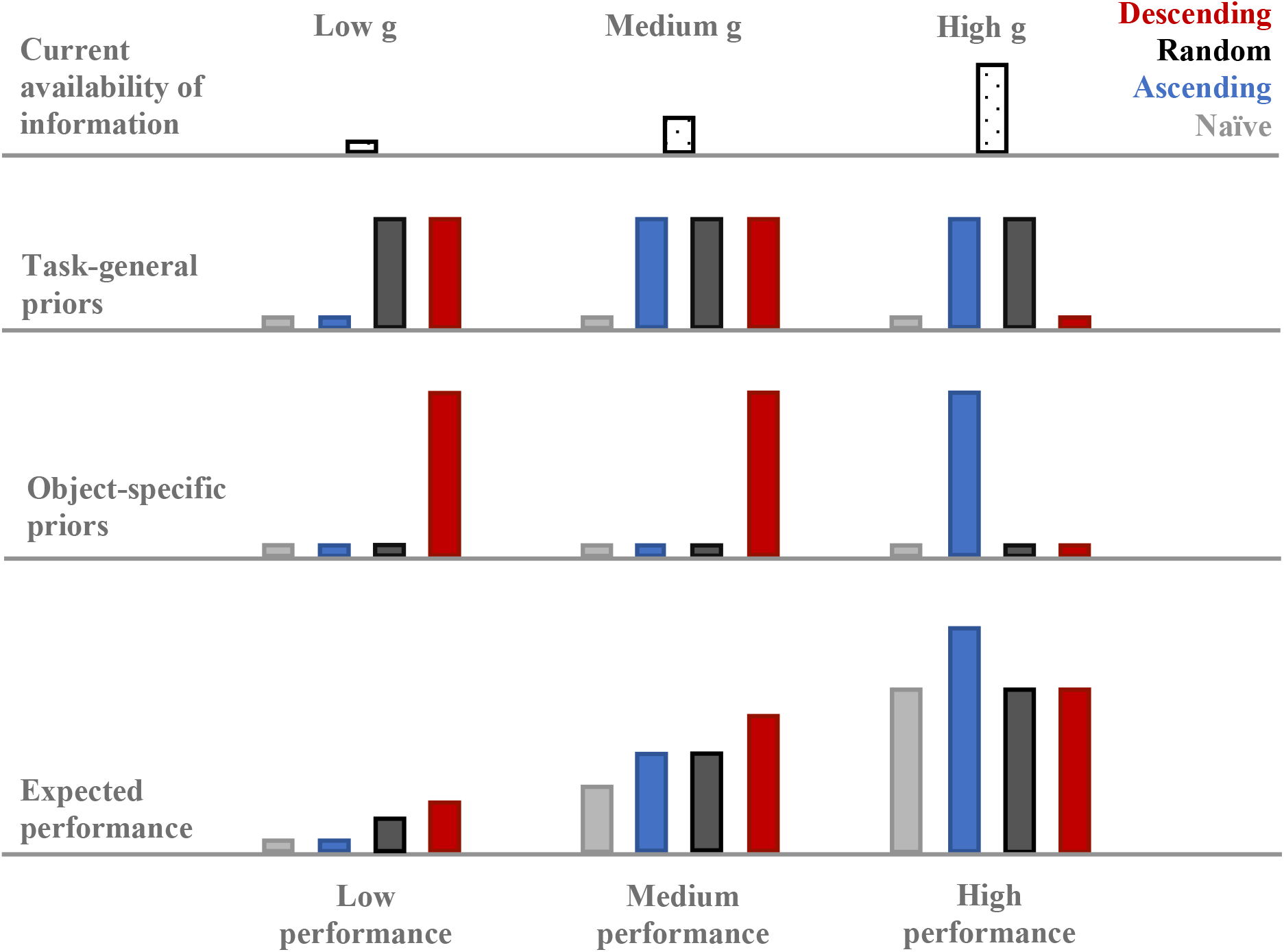
Updated model of participants’ access to information (top row), task-general (middle-top) and object-specific (middle-low) priors, as well as expected performance (bottom) along 3 generic *g*-levels: low (left column), medium (middle column), and high (right column). *Descending* participants are depicted in red, *Ascending* ones in blue, and *Random* ones in black. Grey bars correspond to a hypothetical group of naïve participants building no priors. Higher bars signify higher values.

Follow-up analysis showed that this difference between the behaviour of the two original *Ascending* and *Descending* groups, well predicted by the model, and the unexpected behaviour of the *Random* group, could be related to a strategical difference in the lateralisation of the participants’ gaze. Indeed, we found that all groups compensated the lack of information at low *g* with a higher right lateralisation compared to high *g*, and that the *Random* group overall lateralised more than the other two. This could explain some of the effects for this group’s sampling of less informative points than expected from our model, since the *Random* participants were gazing to the side of the images more than the other participants, while potentially still being able to find informative points to gaze besides them. However, this still translated into a lower recognition performance than the other groups, suggesting that this strategy was not particularly efficient for object recognition.

It is not the first time that randomness is shown to impair behaviour, even when the dimension of randomisation is orthogonal to the task’s goal (e.g., Southwell et al., 2017; Zhao et al., 2013), as is the case here since predicting the *g*-level of the next *Random* trial does not help identifying the object to come. The detection and processing of predictable events has been shown to be both faster and more accurate than unpredictable events (e.g. Correa & Nobre, 2008; Rohenkohl et al., 2012; Bendixen, 2014; Barnes & Jones, 2000). The results for the *Random* group, although diverging from our model’s predictions, are thus compatible with current views on randomness’ effect on learning. Still, in the visual modality, randomness was mostly studied in the context of visual search and had never been shown to impair a process as basic as single object recognition.

It has been suggested that the hindering effects of randomness are due to its low saliency compared to predictable events, resulting in attentional capture and facilitation of the processing of information related to predictable events over unpredictable ones (Turk-Browne et al., 2009; Zhao et al., 2013). This notion was questioned by experiments showing that randomness could impair the processing of information emanating from predictable sources too. Indeed, when Southwell et al. (2017) asked participants to track two concurrent streams of sounds presented binaurally, which could or not be randomised, they showed that whether the target stream was randomised or not did not seem to influence performance per se, and that it was rather the mere tracking of random information that appeared to impair participants’ ability to detect targets in either stream. Indeed, tracking a stream of each type was easier than tracking two random streams, and harder than tracking two regular ones, but finding a target in the regular or the random stream when both were presented was just as difficult. They concluded that neither predictable nor random information seemed to capture attention more, but that the difference between predictable and random information lied in a cognitive load or computational demand disparity. Indeed, processing irregularities is particularly demanding as it constantly generates prediction errors when compared to the observers’ expectations, which requires a constant update of their model, while regular stimuli can be easily explained away by a predictive rule that does not need to be updated with each stimulus, and thus requires less resources.

In our case, only object-specific priors were shown to be impaired, while task-general priors appeared to remain functional: the task’s randomness did not impact all types of information equally, which questions the idea of a general higher attentional capture for regular compared random stimuli (Turk-Browne et al., 2009; Zhao et al., 2013). However, it remains compatible with the hypothesis of an extra cognitive load associated with random events (Southwell et al., 2017). We propose that participants remained able to encode information relative to their building of a general model of their environment (the task) but were unable to store detailed information about specific items of this environment (the objects) while their model of the environment remained hard to build due to its randomness. This appears as an evolutionarily effective behaviour: in a cognitively taxing context when observers are involved in the demanding task of trying to predict items that escape their expectations, encoding items’ specificities in the absence of a general rule for how these items are organised, appears secondary and inefficient. Orhan and Jacobs (Orhan & Jacobs, 2014) proposed that unpredictable stimuli, such as shapes that do not predict colour, provoke a “model mismatch” between participants’ general model of the world from their long-term priors (e.g., bananas are usually yellow) and the information that they are currently experiencing (e.g., a blue banana). We add that the cognitively demanding resolution of this mismatch by updating the model appears to be taking precedence over other types of computations, resulting in participants’ poorer performance in random contexts, linked to reduced specific but not general priors. These results bring important and novel information for the field’s understanding of how randomness impacts perception. They also suggest that perception might be better viewed as a process which is considerably subjective, even at the basic level of object perception as was investigated here.

## 5. Conclusion

Our results indicate that priors already guide basic visual information sampling for a process as fundamental as single object recognition. We show that depending on their priors, participants do not sample the same information: from a more general standpoint, this suggests that priors can cause people to experience the world in fundamentally different ways because at a very basic level, they already sample different pieces of information, possibly coming to different conclusions when integrating them. Furthermore, the general structure through which the information was presented seemed to influence this fundamental process of guiding exploration with priors: randomness, even when introduced in a dimension orthogonal to the task’s goal, destroyed this ability to guide exploration through specific but not general priors. Participants in the randomised paradigm behaved seemingly naïvely despite their intermediate access to information. These findings have important societal implications for how we structure information in everyday situations, such as teaching or policy making, and stress the importance of catering for different stages of learning and presenting information in a structured manner.

## 6. Materials and Methods

This study was based on work from Moca et al. (2011). They recorded the data and analysed two of the three groups of participants (the *Ascending* and *Descending* groups, leaving the *Random* group’s data unexplored). Here, the data was entirely re-analysed, including the analysis of the yet unreported *Random* group, which this paper focuses on.

### 6.1. Participants

18 participants (10 females, mean age 28.3 years, S.D. 4.4 years) took part in this experiment. They either joined as volunteers or received course credits for their contribution as part of their undergraduate Psychology curriculum. They all had normal or corrected-to-normal vision. They each were assigned to one of three experimental conditions (described below and referred to as *Ascending*, *Descending*, or *Random*), resulting in three groups of N=6 subjects each. All participants gave their written consent before starting the experiment. All procedures were approved by the local ethics committee of the University of Medicine and Pharmacy “Iuliu Hatieganu” of Cluj-Napoca, Romania, under the approval No. 150/10.12.2009.

### 6.2. Stimuli

Moca et al.’s (2011) stimuli were used: images were generated via the *Dots* methods, through which contour information extracted from source images of objects was used to apply a deformation to a 2D lattice of dots to displace dots and reveal objects’ outline. The visibility of the stimuli and their information content was precisely manipulated through the use of 7 different deformation levels controlled by a gravitation constant *g*. It ranged from *g* = 0.00 at the lowest visibility level (no deformation) to *g* = 0.30 at the highest visibility level, with steps of *g* = 0.05. For each of the 50 objects, 7 stimuli at each of the 7 *g*-levels were generated, resulting in a pool of 350 stimuli. Example stimuli for one object are shown in **Figure 12**.

**Figure 12.**
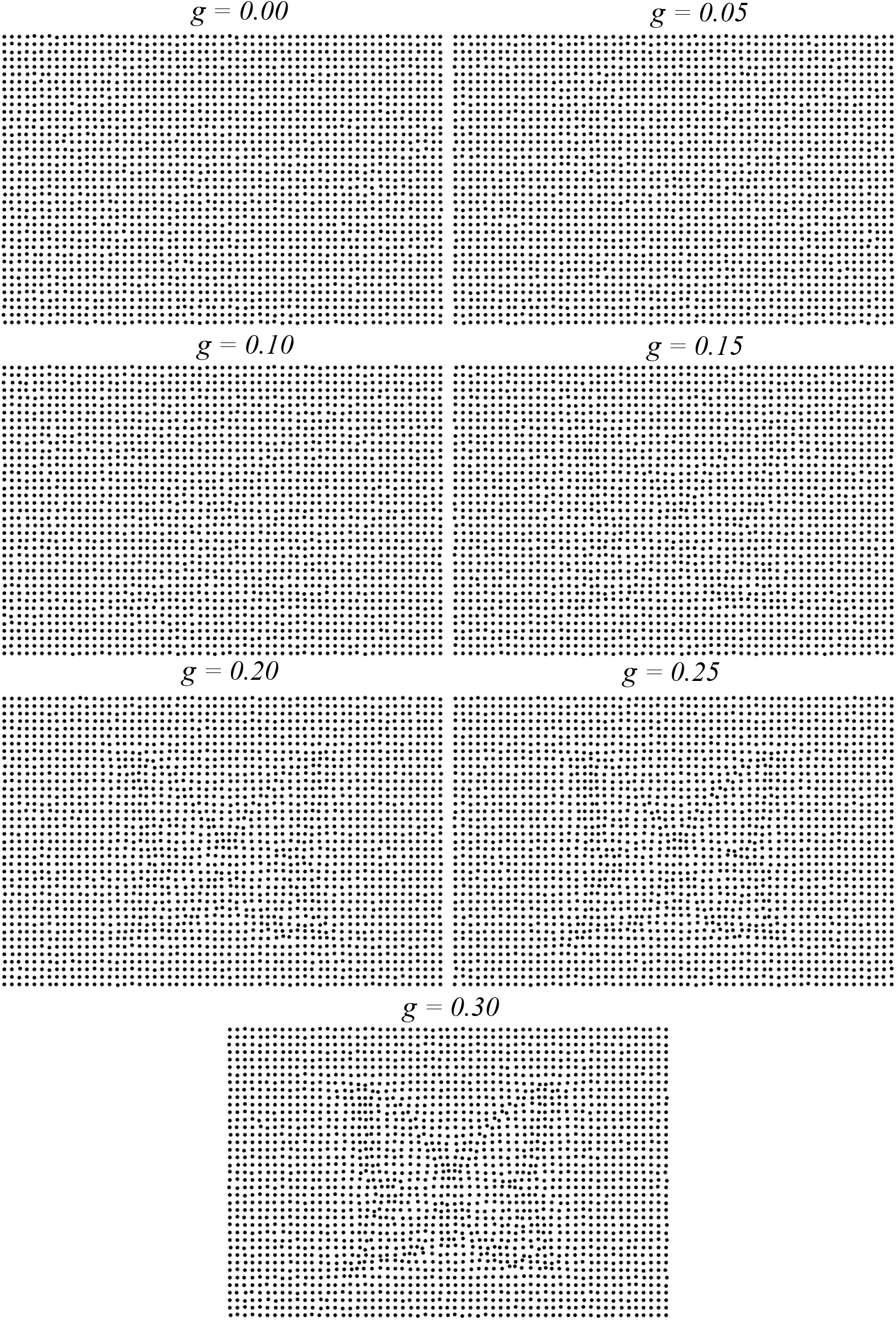
Example of *Dots* stimuli for one object at each level of visibility *g*, shown in order of increasing *g*.

### 6.3. Procedure

Participants were tested in a detection-recognition task. Stimuli were presented on a 22” Samsung SyncMaster 226BW (2ms Grey-To-Grey response time) at a resolution of 1,680 x 1,050 pixels, placed at a distance of 115cm from the participant. The images were presented in the central part of the screen at a resolution of 600 x 400 pixels. Each trial started with a red fixation cross for a random duration of 1,000 to 1,500ms. The stimulus then appeared on the screen for an indefinite duration. Participants were instructed to visually explore each stimulus for as long as they wanted to and to decide whether the dots pattern of the stimulus represented anything meaningful. After visualising the stimulus and when they wanted to, they pressed one of three buttons to indicate whether they had “*seen*” the object (“L” key: they perceived something meaningful in the stimulus and knew what it was), were “*uncertain*” what the object was (“S” key: they thought they saw something but were not sure what it was), or saw “*nothing*” in the dots grid (“A” key: they did not think anything meaningful was depicted). This was followed by a green fixation cross for 500ms, after which a message appeared asking the subject to explicitly name the object that they (thought they) had seen, in the cases when they answered “*seen*” or “*uncertain*” (guess). Their oral answers were manually recorded by an experimenter present in the room throughout the experiment (only “*seen*” answers were considered for the calculation of the accuracy measure). Participants finally pressed SPACE to move on to the next trial, which began after a 200ms delay. The session started with a 14-trial training block using a separate set of objects, followed by the experiment’s blocks. The general design for the trials is summarised in **Figure 13**.

**Figure 13.**
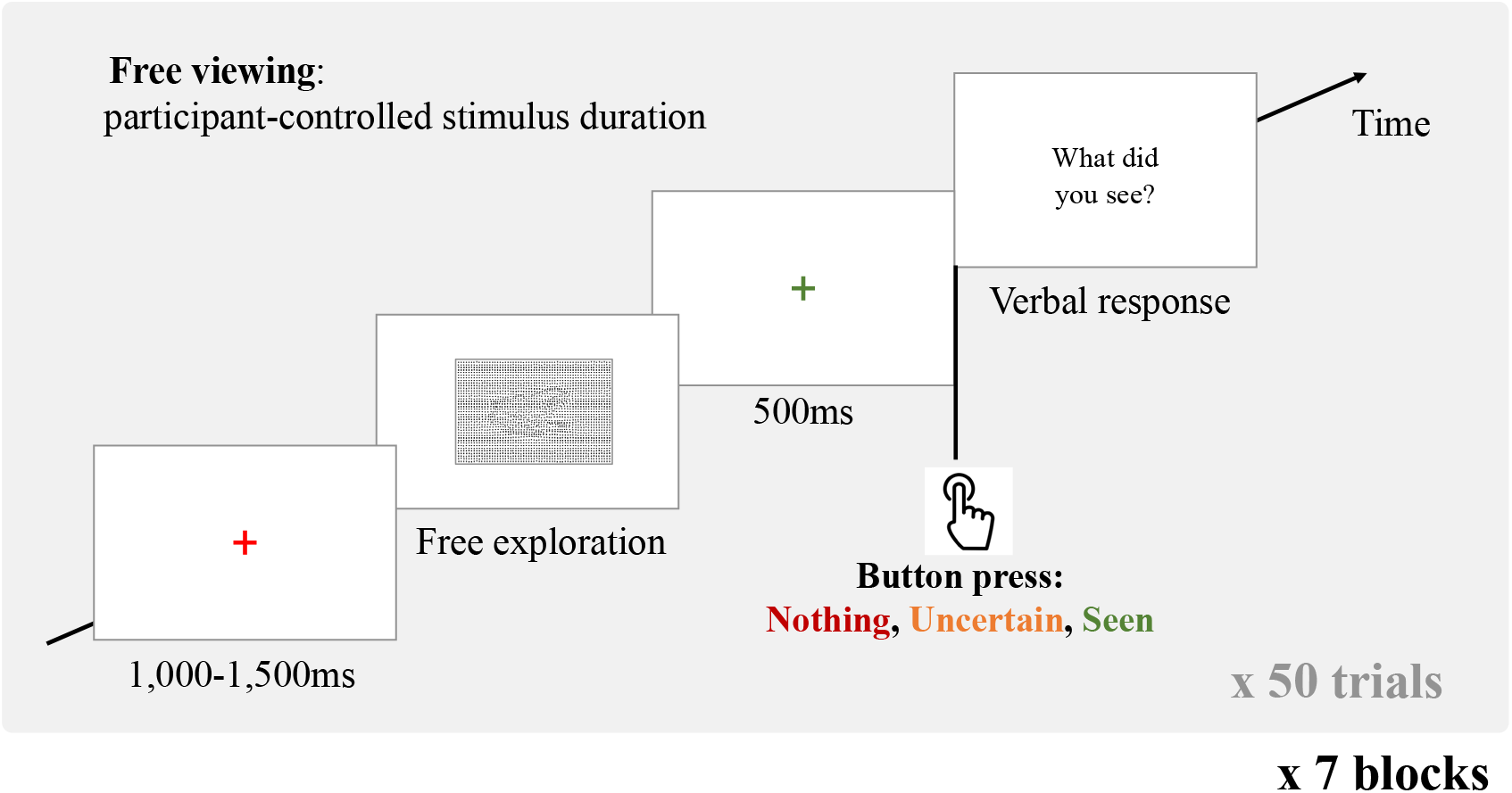
Structure of a trial; the trials were composed of a fixation cross of randomised duration in the 1,000-1,500ms interval, a free viewing exploration phase until the participant pressed one of three buttons to signify that they had viewed “Nothing”, were “Uncertain” or had “Seen” an object, after what they were prompted by a sentence on the screen to verbalise the object they thought that they saw if the pressed “Uncertain” or “seen”. There were seven blocks of 50 trials, and each object was shown one per block, at each of the seven *g*-levels across all blocks.

### 6.4. Presentation order – between-subjects design

Stimuli were presented in 7 blocks, each containing all 50 objects at one of the 7 visibility levels. The order of the objects within the blocks was randomised for each participant. The order of the objects’ visibility throughout blocks varied between the groups to manipulate the participants’ access to information. One group of participants saw the stimuli in an *Ascending* fashion: the stimuli were presented at the same visibility level within blocks, starting with a block of the lowest (no deformation) visibility level, going up to more and more visible stimuli at each block. A second group viewed the stimuli in a reversed *Descending* path, where they first saw the objects at the highest visibility level, going down in visibility at each block. Both these groups correspond to those described in Moca et al. (2011). Finally, a last group viewed the stimuli in blocks of mixed visibility levels, in which all the objects were presented once per block, but each at a *Random* level of visibility out of the seven. This latter presentation order resulted in an access to visual information which was intermediate between the *Ascending* presentation (lowest, uninformative content first) and the *Descending* presentation (highest, most informative content first). This group was a new addition to the data presented by Moca et al. (2011).

### 6.5. Recordings

Participants’ button-presses (Seen/Uncertain/Nothing) were recorded with precise timings, together with their verbal responses (object’s name). Eye-tracking was used to monitor their gaze throughout the experiment. The eye-tracking recordings were made monocularly with an ASL EyeStart 6000 system at a rate of 50Hz. Participants’ heads rested on a cheek-rest to avoid changes in their head position during the tracking, while still enabling them to speak after each stimulus. A nine-point calibration was conducted at the start of each block, and each trials’ fixation cross was used as a post-hoc calibration to correct for potential shifts in the eye position within blocks.

### 6.6. Data processing

#### 6.6.1. Manual inspection of the data vs. Moca et al.’s automatic pipeline

Although two of the groups were already presented in Moca et al. (2011), all datasets from all three groups were processed anew from raw data for the present study. The same pre-processing pipeline was used, which included automatic identification of saccades and fixations, but all fixations were then manually checked for any missing, additional, or misidentified saccades. This was found to increase the quality of the data, as shown by a comparison of fixation durations between the automatic process in Moca et al.’s data and the current manually checked data. A histogram of the fixation durations, pooling all trials of all *Ascending* and *Descending* participants, for the fully automatic process (**Figure 14A**) and for the manually checked data (**Figure 14B**) showed that manually checking the data resulted in an overall more normal distribution, with less of the very short and very long fixations, and mean and median values closer to each-other (the time bin for the mean and median values in the automatic pipeline were, respectively, 480-500ms and 900-910ms, which shifted to 540-560ms and 800-820ms after manual inspection). There were no substantial changes in the results between the manual and the automatic processing, thus the difference between the pre-processing pipelines is not further discussed in the paper.

**Figure 14.**
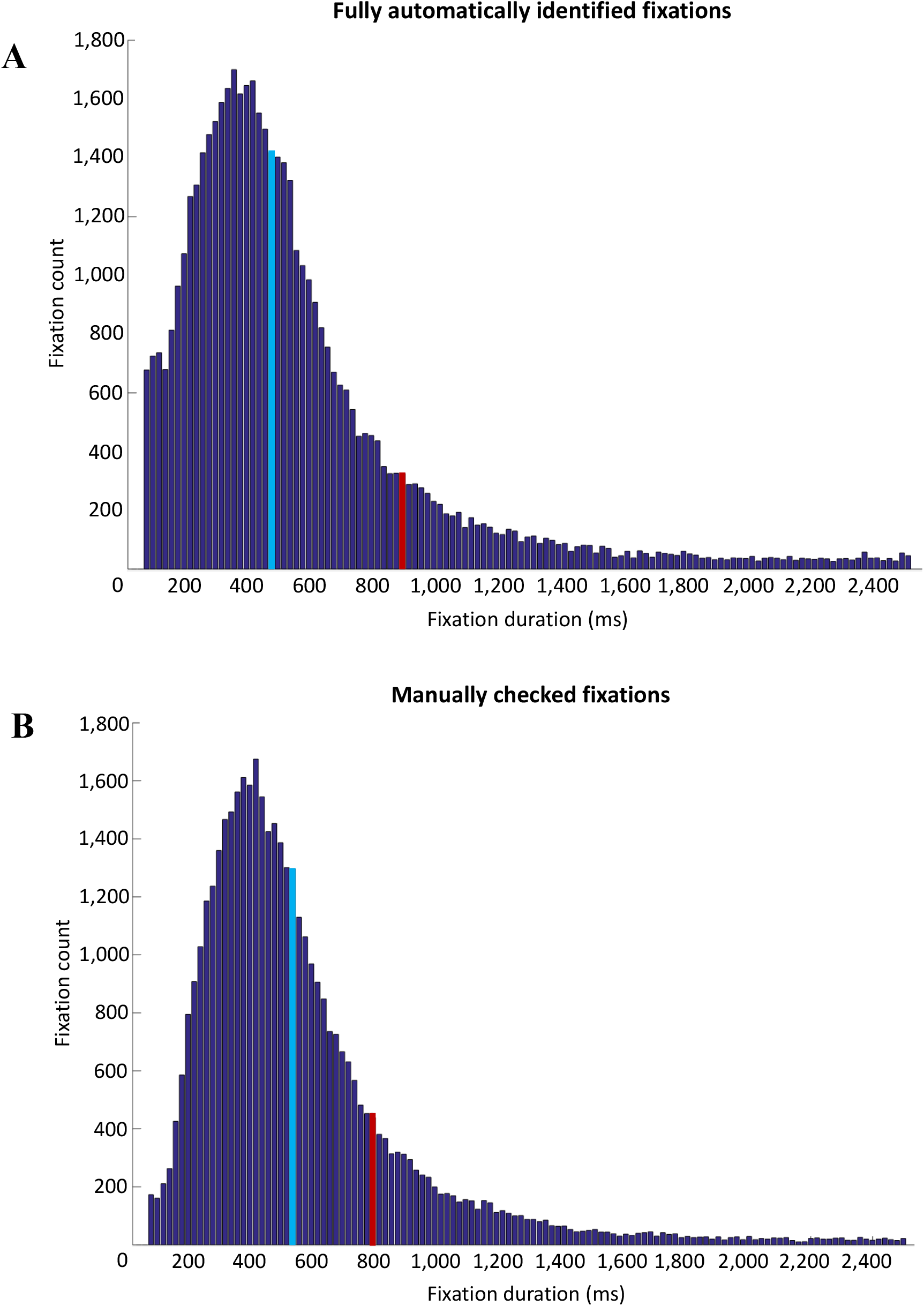
Histogram, using 20-ms time bins, of the fixation durations of all *Ascending* and *Descending* participants across all trials for the fully automatic process (A) and the manually checked data (B). The blue bin represents the bin that contains the median, and the red bin, the mean. The tail of histogram was too long for good visualisation and was not entirely plotted here.

#### 6.2.2. Pre-processing pipeline

Trials were automatically screened and any trial with over 50% data loss was discarded. Horizontal and vertical gaze location data were smoothed, and fixations were automatically detected using a simplified version of Nystrom and Holmqvist (2010) algorithm with two adaptive velocity thresholds: the velocity of the eye’s position was computed and summed to obtain a horizontal-vertical composite velocity variable. Each time the velocity crossed a high threshold, a saccade was detected, whose start and end were identified using a second, lower threshold. The thresholds were computed on a trial-by-trial basis according to the trial’s mean and standard deviation in the composite velocity, respectively 4 and 1.5 standard deviations above the mean. Fixations were defined as a time series between two saccades with a minimum duration of 50ms. This automatic algorithm was used to guide the parsing of the data but, as described above, a manual check of all saccades for all trials and all participants was conducted to avoid misdetections of saccades in case of local noise. The first and last fixations were discarded as they presented a significantly different profiles in standard eye-movement statistics (duration, spread) and were linked to, respectively, base-line fixations on a central cross and post-button click awaiting fixations before verbalising the answer (Henderson, 1993).

At each fixation identified, the stimulus’ local information content around this point was reconstructed in two different ways. On the one hand, the information *physically* conveyed by the image was reconstructed from the stimulus image itself: taking an area of 0.5 visual degrees around the fixation, we calculated the amount of *local dots displacement (LDD)*, which directly relates to the information present at this location. On the other hand, we also calculated a more semantic*, hidden* form of local information content, using the object’s source image this time: in the same area of 0.5 deg. around the fixation, we calculated the amount of *local contour density (LCD)* in the source image. Since the squared contour density was used to generate dots displacement in the stimulus, these two measures are highly related, however they do not fully map onto each other.

## 7. Acknowledgements

The authors wish to thank the participants who took part in this study, the researchers who helped with the data collection and processing, notably Cristina Pelea, and those who further improved this project through inspiring discussions and comments. This work was funded by a European Union Horizon 2020 Research and Innovation Marie Skłodowska-Curie doctoral grant No 721895, a postdoctoral Wellcome Trust Institutional Strategic Support Fund grant distributed by Birkbeck, University of London, a NO (Norway) Grant 2014-2021, under project contract number 20/2020 (RO-NO-2019-0504), four grants from the Romanian National Authority for Scientific Research and Innovation, CNCS-UEFISCDI (codes PN-III-P3-3.6-H2020-2020-0109, ERA-NET-FLAG-ERA-ModelDXConsciousness, ERANET-NEURON-UnscrAMBLY, and GAMMA-CXCD code PN-III-P1-1.1-TE-2021-0709), and a H2020 grant funded by the European Commission (grant agreement 952096, NEUROTWIN).

## 8. Competing interests

All the authors declare no competing interests.

## Notes

### Competing Interest Statement

The authors have declared no competing interest.

### Summary of Updates

Changed Figures 9 and 10, and revised the text that describes lateralisation results. Also, we added some more details to the discussion section.

